# High-throughput measurements of intra-cellular and secreted cytokine from single spheroids using anchored microfluidic droplets

**DOI:** 10.1101/2020.09.25.312900

**Authors:** Adrien Saint-Sardos, Sebastien Sart, Kevin Lippera, Elodie Brient-Litzler, Sebastien Michelin, Gabriel Amselem, Charles N. Baroud

## Abstract

While many single-cell approaches have been developed to measure secretion from anchorage-independent cells, these protocols cannot be applied to adherent cells, especially when these cells requires to be cultured in 3D formats. Here we demonstrate a platform to measure the secretions from individual spheroids of human mesenchymal stem cells, cultured within microfluidic droplets. The platform allows us to quantify the secretion from hundreds of individual spheroids in each device, by using a secondary droplet to bring functionalized micro-beads into proximity with each spheroid. We focus on vascular endothelial growth factor (VEGF) and measure a distribution of secretion levels that presents broad heterogeneity within the population of spheroids. Moreover, the intra-cellular level of VEGF-A on each spheroid, measured through immuno-staining, correlates well with the extra-cellular measurement, indicating that the heterogeneities observed at the spheroids level result from variations at the scale of individual cells. Finally, we model the molecular accumulation within the droplets and find that physical confinement is crucial for measurements of protein secretions. The model predicts the time to achieve a measurement, which scales with droplet volume. Therefore these first measurements of secretions from individual spheroids provide several new biological insights.

## I. INTRODUCTION

A wide range of secreted molecules such as hormones, neurotransmitters, or cytokines, provide a vehicle for cell-cell communication and help regulate the function on the scale of whole organs and organism. The different types of molecules are present at different concentrations *in vivo*, and act over a range of time scales, from seconds to days. Indeed, variations in the rate of secretion of different molecules indicate the state of the cells, for example as a response to changes in their environment. Standard methods for measuring secreted molecules in typical cell culture experiments however require the presence of a large number of molecules, which translates into volumes of tens to hundreds of *µ*L and give information at the level of large populations of cells. As a result, these techniques hide the heterogeneities that may exist on the scale of individual cells. It is becoming increasingly clear, however, that these heterogeneities are present and play a determining role in many biological processes, such as the immune response to an unknown pathogen [1], or as a prognostic marker for some cancer types [2].

For this reason, a wide range of methods have been developed for quantifying the secretions of individual cells, in particular by using microfluidics [3]. These methods always rely on reducing the volume to be analysed to become close to the scale of a cell. One early approach involved encapsulating the cells in hydrogel [4, 5], which enabled their quantification by flow cytometry. More recently, microfabricated wells were used to isolate individual cells and surface functionalization was used to selectively capture specific secreted molecules [6]. This method was further developed for the quantification of the heterogeneity in secretion of cytokines by immune cells [7]. Alternatively, the functionalized solid can be the surface of a micro-bead, which can be co-incubated with the cells. Such beads can be located within the microfabricated chambers [8, 9], or they can be placed downstream of the cells in a mean flow [10]. finally, droplet microfluidics has also been used to co-encapsulate cells and functionalized beads within droplets [11–14], thus taking advantage of all of the features of droplet microfluidics, including simplified microfabrications, encapsulation, as well as the different tools existing for droplets [15].

The efforts to measure single-cell secretions however have focused almost exclusively on non-adherent cells, which represent a small fraction of all secreting cells in the body. In contrast, the case of anchorage-dependent cells is more complicated for several reasons. First, these cells must adhere to a solid substrate in order to survive, making many of the existing microfluidic technologies difficult to use. Second, the culture format often alters the biological activities of these cells. Indeed, it has been shown that 3D culture formats better promote the biological functions of cells, including their secretory activities [16–19]. Third, the cell’s phenotype, and by extension the molecules it secretes, depends on its local microenvironment. This implies that the behavior *in vivo* is not a simple superposition of the behavior of individual cells in isolation; instead, the cells can organize into functional units whose secretions are determined by the interactions between them, as it can be exemplified by the maturation of liver functions that is mediated by a complex interplay between hepatic, endothelial and stromal cells [20]

The challenge of 3D culture can be addressed by using spheroids, which emerge from the clustering of hundreds of adherent cells into a coherent functional cellular unit. When originating from a heterogeneous population, the cells in the spheroid self-arrange in an organized spatial manner, such that the cellular function is linked with the 3D structure [21]. As such they constitute an *in vitro* system that recapitulates some of the complexity of the *in vivo* conditions, with a relatively simple production protocol [22]. However, it is not known how the heterogeneities at the single cell level translate at the spheroid scale. As such, the heterogeneity of the secretion at the spheroid scale is arguably more relevant to understand *in vivo* biological behavior than single-cell measurements. Consequently, there is a need for platforms for the formation and culture of spheroids, while enabling high-throughput quantitative information on individual spheroids and their secretions.

Here we quantify the secretion of vascular endothelial growth factor (VEGF-A), a proangiogenic molecule [23], by spheroids made with human mesenchymal stem cells (hMSCs). hMSCs constitute a heterogeneous population of progenitors of several cellular types of connective tissues that shows important trophic functions while cultivated in 3D [24]. To interrogate the heterogeneity of protein secretion at the single spheroid level, we use an integrated droplet microfluidic device, which is equipped with capillary anchors that contain two trapping areas [25]. This trap geometry allows it to be divided into a culture area and an analysis area, to which the tools to immobilize and to detect secreted proteins can be brought at any desired time point during the culture period. Through quantitative fluorescent imaging, the platform enables the combined culture of dense arrays of spheroids, their characterization, as well as the quantification of their secretions *in-situ*.

## II.MATERIALS AND METHODS

### a. Microfabrication

The microfluidic chips were fabricated using the protocols detailed in our previous studies [17, 21, 25]. Briefly, the chip consists in a top part made with Poly(dimethylsiloxane) (PDMS, SYLGARD 184, Dow Corning, 1:10 (w/w) ratio of curing agent to bulk material) on which is inprinted a flow focusing device connected to an emulsification channel (serpentine), diverging rails, and terminated by a culture chamber; the bottom part of the chip is a PDMS layer (about 400 *µ*m thick) etched with an array of capillary traps, which is bounded on a microscopy glass slide (Figure 1A). The top and bottom parts of the chip were assembled after plasma treatment (Harrick, Ithaca, USA). The chips were rendered fluorophilic by filling them three times with Novec Surface Modifier 1720 (3M, Paris, France), a fluoropolymer coating agent, for 30 min at 110°C on a hot plate.

**Figure 1.**
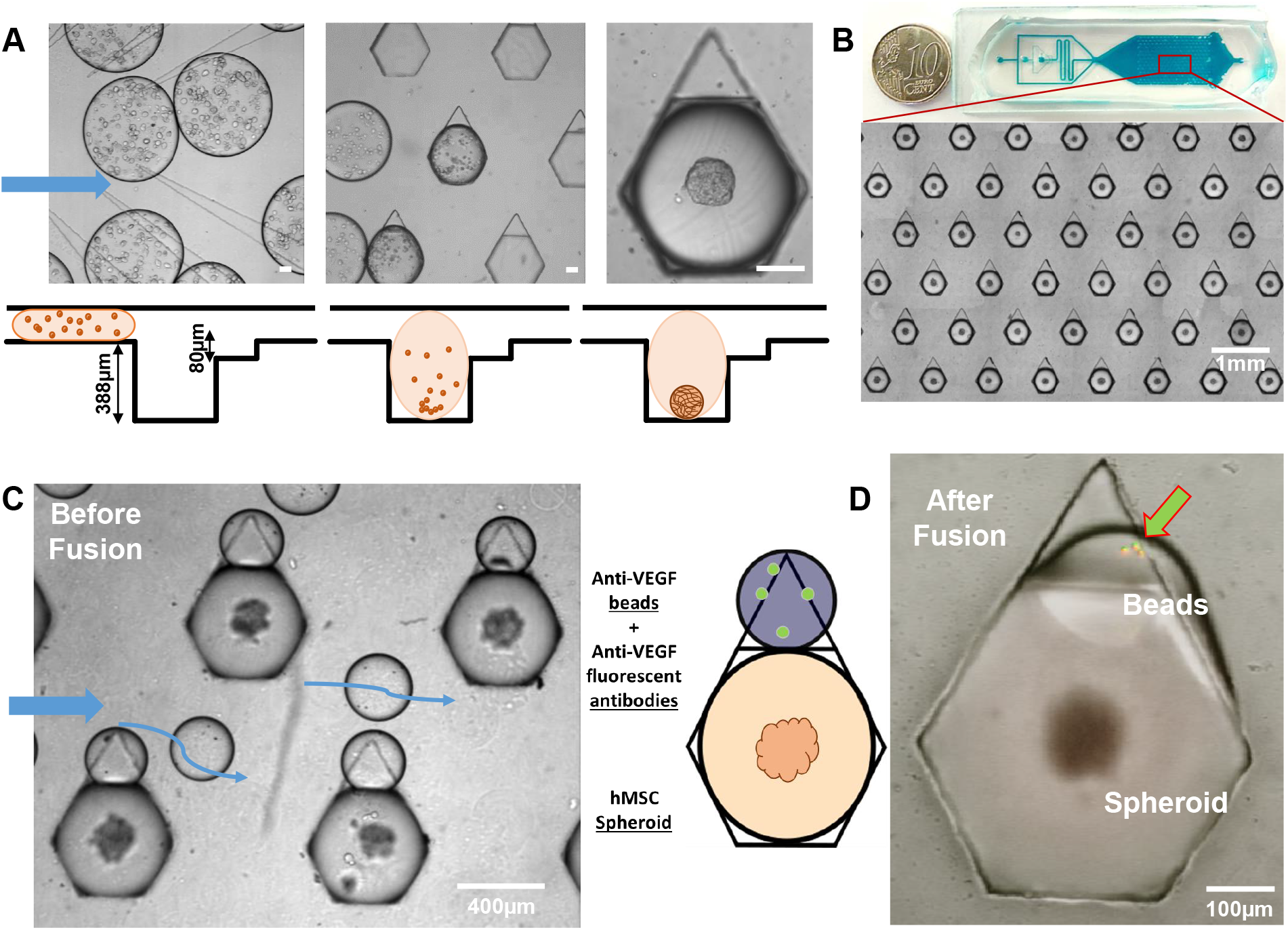
Microfluidic platform for spheroid formation and VEGF-A quantification. (A) Top view (top row) and side view (bottom row) of the microfluidic loading process: cells are loaded as free suspensions in liquid agarose droplets (left) that get trapped in the hexagonal capillary anchors (middle). Over the next 4 hours, cells sediment and form 3D spheroids (right). Scale bars: 100 *µ*m. (B) Top view of the microfluidic culture chamber: the microchip contains an array of 18×14 spheroids of hMSC on a surface of 2.4 cm^2^. (C) Top view of the secondary droplet trapping next to spheroid-containing drops: anti-VEGF-A beads and anti-VEGF-A secondary fluorescent antibody are also introduced into liquid agarose droplets and get trapped in the smaller anchor next to the spheroid droplet. (D) Top view of the gelled agarose droplet 12 hours after droplet fusion: beads are visibly fluorescent due to their barcode (green) and their capture of VEGF-A (red).

### b. Cell Culture

Human mesenchymal stem cells derived from the Wharton’s Jelly of umbilical cord (hMSCs) (ATCC PCS-500–010, American Type Culture Collection, LGC, Molsheim, France) were obtained at passage 2. hMSCs were cultivated in *α*-MEM (Gibco, Life Technologies, Saint Aubin, France) supplemented with 10% (v/v) fetal bovine serum (Gibco) and 1% (v/v) penicilinstreptamicine (Gibco), as previously described [21]. The cell population was positive for CD73, CD90, CD105, and CD146, but not for CD31, CD34, CD14 [21]. In addition, hMSCs were capable to be differentiated towards osteoblastic, chondrogenic and adipogenic lineages [21].

### c. Spheroid Culture

The formation of spheroids with hMSCs was performed as previously described [21]. Briefly, a 50 *µ*L solution of 4.10^6^ cells/mL in 0.9% (w/v) liquid agarose was loaded into a 100 *µ*L glass syringe (SGE, Analytical Science, France), while Fluorinert FC-40 oil (3 M, Paris, France) containing 2%(w/w) PEG-di-Krytox surfactant (RAN Biotechnologies, Bervely, USA) was loaded into 2.5 mL glass syringes (SGE, Analytical Science). Droplets of cell-liquid agarose (of about 50 nL) were formed at the flow focusing junction, by controlling the flow rates using syringe pumps (neMESYS Low Pressure Syringe Pump, Cetoni GmbH, Korbussen, Germany). After complete loading, the chips were immersed in PBS and the cells were allowed to settle down and to organize as spheroids for 4 h in a CO2 incubator. Then, the agarose was gelled at 4°C for 30 min [17, 21].

### d. Viability Assay and Immunolabelling of Intracellular VEGF-A on Chip

All the steps for fluorescent staining were performed on-chip [17, 21]. The cell viability was assessed using a ReadyProbes^™^Cell Viability Imaging Kit (Blue/Red), (ThermoFischer), as previously described [17, 21]. The intracellular production of VEGF-A was measured by immunocytochemistry, following the protocols detailed in [21]. Briefly, the cells were fixed in a 4% PFA, then permeabilized with 0.2-0.5% (v/v) Triton X-100 (Sigma-Aldrich). The samples were blocked with 5% (v/v) FBS in PBS and incubated with a solution of rabbit anti-human VEGF-A monoclonal antibody (ab52917, Abcam, Cambridge, UK) diluted at 1:100, and then revealed using the Alexa Fluor ^®^ 647 conjugate goat polyclonal anti-rabbit IgG secondary antibody (711-605-152, Jackson Immunoresearch Europe Ltd, Cambridge, UK) diluted at 1:100 in 1% (v/v) FBS, for 90 min. Finally, the cells were counterstained with 0.2 *µ*M DAPI (Sigma-Aldrich).

### e. Immunodetection of Extracellular VEGF-A on Chip

The micrometric beads coupled with capture antibodies and the biotinylated-detection antibodies cocktails against human VEGF-A were prepared and diluted following the manufacturer instructions (Luminex Corporation eBioscience, Austion, Texas, US). Then, the biotinylated-detection antibodies cocktail was mixed with a diluted streptavin-phospj solution (eBioscience, Inc., Thermo Fisher, Waltham, Massachusetts, USA), following the manufacturer instructions. Droplets of volume *V*_*d*_ ≈ 50 nL containing a solution of 3% agarose as well as cells or a serially diluted recombinant VEGF-A solution (Invitrogen, Carlsbad, California, USA) were first immobilized in the larger trapping area of anchors. On the day of the experiment, a smaller drop of volume ≈ 5 nL containing 4-5 beads coupled with the capture antibodies, the detection antibodies conjugated with phycoerythrin (PE) and a 3% agarose solution were trapped in the smaller area of the traps. The small and large drops were fuses by flushing a 50% PerFluorOctanol (PFO) solution diluted in NOVEC HFE-7500 (3M, Paris, France), which was then washed away by flushing pure FC-40 into the microfluidic device. Ten hours post droplet fusion, the fluorescent signals of the beads were acquired under a motorized wide-field fluorescent microscope (Nikon, France), using a cooled-CCD camera (ANDOR, Oxford Instruments, Abingdon-on-Thames, UK) and a 10X-long working distance objective (Nikon, France). The calibration curve linking the VEGF-A concentration *C* in the droplet with the measured fluorescence *F* is fitted by a pseudo-sigmoidal model[26], using four constants *b*_*i*_:

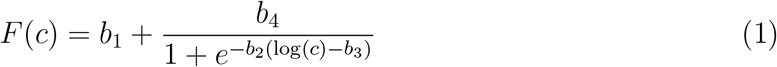

### f. Image, Analytical and Statistical Analysis

Each experiment was repeated three times (N= 3 chips per conditions). Image analysis, data treatments, and statistical analysis were performed using ImageJ [27] and Matlab 2017b (The MathWorks, MA, USA). We used bandpass filtering on the fluorescence images to create binary masks where only the beads are visible, enabling to obtain the pixel values of the VEGF-A-related fluorescence at the precise bead positions. This fluorescence signal was averaged over each bead, and the local fluorescence background was subtracted from the bead signal. Significance testing was performed over the spheroid population with the Wilcoxon-Mann-Whitney test for non-normal distributions. *: p < 0.05; **: p < 0.01; ***: p < 0.001; N.S.: non-significant. In theses tests, each spheroid is considered as a single biological replicate.

### g. Smoldyn stochastic simulations

We use the software Smoldyn (http://www.smoldyn.org) to model VEGF-A secretion, diffusion and capture within the droplet. Smoldyn performs stochastic spatial simulations by treating molecules as point-like particles that diffuse and react, using a framework introduced by von Smoluchowski [28]. Our Smoldyn simulation files are available in the Supplementary Material. Briefly, we simulate a spheroid of radius *R*_*s*_ = 40 *µ*m centered in a droplet of radius *R*_*d*_ = 200 *µ*m. The spheroid secretes VEGF-A at a rate A = 20 molecules*/*s, and VEGF-A then diffuses in the droplet with a diffusion coefficient *D* = 133 *µ*m^2^*/*s. A capture bead of radius *a* = 5 *µ*m is placed in the droplet, 7.5 *µ*m away from the droplet border. In the experimental system, the number of binding sites on the capture bead is in large excess compared to the number of VEGF-A molecules secreted by the spheroid, and binding of VEGF-A to the bead is much faster than secretion or diffusion (see section sec:theory). To keep simulation times reasonable, the capture bead is simulated as a perfectly absorbing sphere, and the binding reaction is considered to be diffusion-limited, so that the binding rate is *k* = 4*πDa* [29]. The simulation time step is *t* = 0.01 s.

## III.RESULTS

### A. Platform for spheroid formation and secretion quantification

The protocol takes place in a microfluidic chip equipped with a combined flow-focusing and step junction for the formation of monodisperse water-in-oil microdroplets [17] of volume *V*_*d*_ ≈ 50 nL. The aqueous droplets contain a suspension of hMSCs, culture medium and liquid agarose, and are initially transported in channels whose height is smaller than the droplet diameter. The droplets are then distributed evenly into the culture chamber through guiding rails [30] (Figure 1A left). The floor of the chamber is patterned with asymmetric capillary anchors, that contain a deep and large trapping area (400 *µ*m diameter and 388 *µ*m deep), terminated with a triangular extension of a smaller volume (200 *µ*m long and 80 *µ*m deep), see Fig. 1A. The droplets are firmly anchored in the larger part of the traps, where they adopt a spherical shape, thus reducing significantly their surface energy [31]. The secondary traps anchor less efficiently the drops and remain empty (Figure 1A).

After droplet anchoring, the oil flow is stopped and the cells are allowed to settle down at the bottom of the droplets. This initiates cellular clustering, and the cells spontaneously self-arrange to form spheroids (see Figure 1A right) [17, 21]. The characteristic time for spheroid formation with hMSCs is about 4 hours (Supp. Fig. S1). The protocol results in the formation of 252 spheroids with an average diameter of 100 *µ*m (Figure 1B), that remain viable for at least a 2-day culture period [21].

To measure the level of protein secretion by the spheroids, smaller droplets of volume ≈ 5 nL containing on average 5 magnetic beads grafted with a anti-VEGF-A capture antibody, a solution of anti-VEGF-A detection antibody tagged with the fluorophore phycoerythrin (PE), and liquid agarose at a concentration 3% are loaded into the culture chamber (Figure 1C). These droplets are immobilized in the secondary, triangular, anchors. The two drops containing agarose are then gelled at 4°C, which firmly retains in place the spheroid and the antibody-grafted beads. Both droplets are fused by destabilizing their interface using PerFluoroOctanol (PFO). After a day of incubation, the beads are detected using their barcode signal (in green in Figure 1D), and the fluorescence levels associated to VEGF-A production are quantified by fluorescence microscopy and image analysis.

### B. Calibration of the bead-based protein quantification assay on chip

We begin by calibrating the commercial immunoassay used throughout this work (LH-SCM293, Luminex Corporation, Texas, US) and determine the range of concentrations detectable by the microfluidic method. Drops containing concentrations of VEGF-A ranging from 1.5 × 10^1^ to 2.5 × 10^5^ pg/ml are first loaded in the larger trapping area of the anchors. Then, smaller drops containing on average 5 anti-VEGF-A capture beads as well as detection antibodies are trapped in the triangular extension of the anchor, see Figure 2A. The two droplets are fused as described above (Figure 2B).

**Figure 2.**
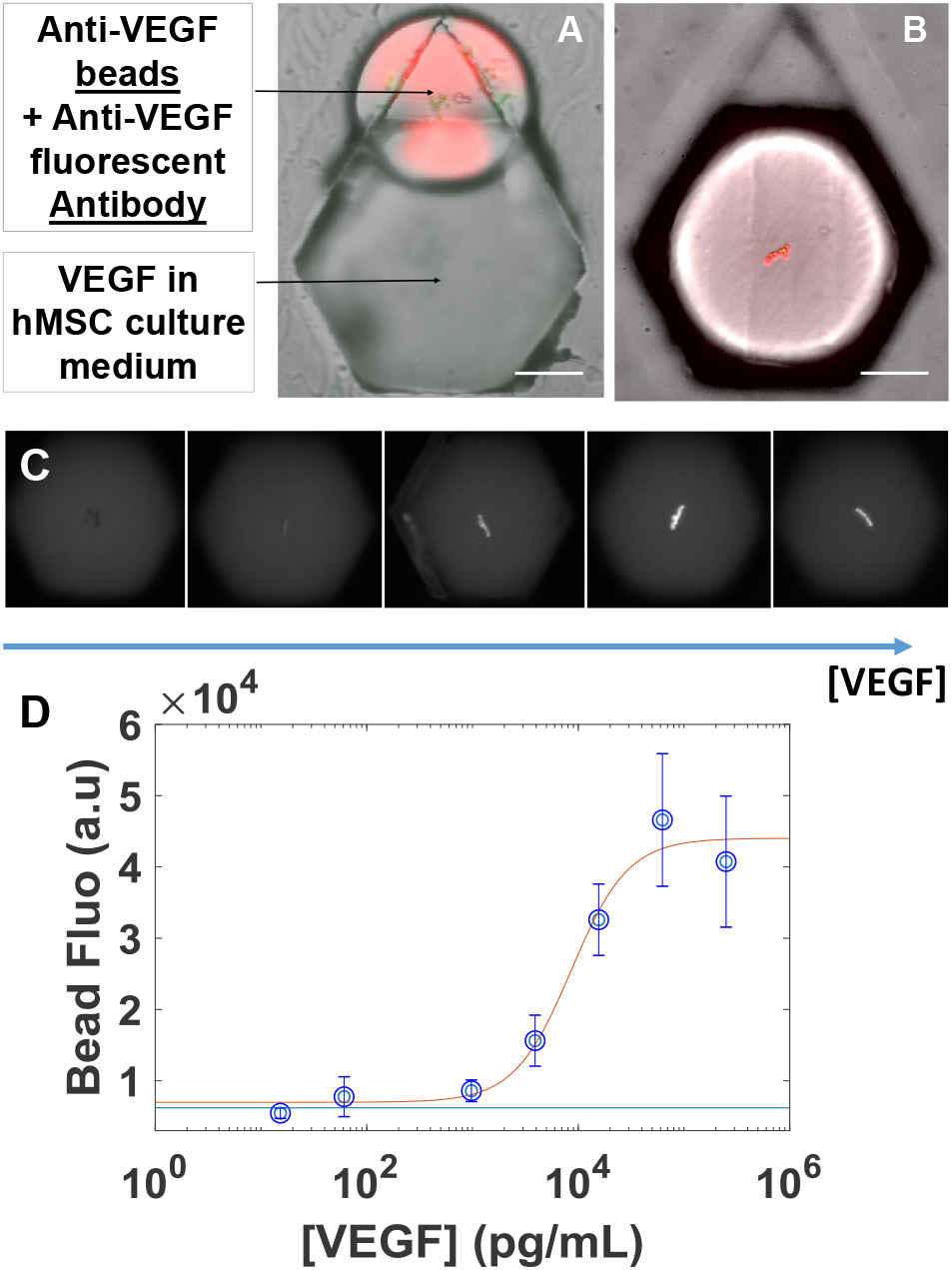
Calibration of the bead immuno-assay. (A) Top view of the experimental set-up for calibration of VEGF-A measurements: the bigger droplet contains VEGF-A at known concentrations between 10^1^ and 10^6^ pg/ml, the smaller droplet contains anti-VEGF-A beads, and anti-VEGF-A fluorescent antibodies. (B) After fusion and 15 hours of incubation, snapshot of beads reacting with 3.10^5^ pg/ml of VEGF-A. In (A) and (B), snapshots combine bright field and fluorescence signals. (C) Typical fluorescence snapshots of the beads after incubation with (0, 1.95, 7.81, 31.25, 125.0, 500.0).10^4^ pg/ml. (D) Mean fluorescence of the beads as a function of the initial VEGF-A concentration in the large drop. n>70 traps per concentration. Bottom blue line displays the mean bead fluorescence measured for [VEGF-A]=0 pg/ml. 4-Parameter Logistic fitting (see Eq.(1)), with parameters *b*_1_ = 6.98 × 10^3^, *b*_2_ = 3.72, *b*_3_ = 3.93, and *b*_4_ = 3.70 × 10^4^.

To ensure that steady-state has been reached and all VEGF-A molecules have bound to the bead, the assay is incubated for 15 hours, a time much longer than the typical times associated with diffusion (see Section III E for a detailed discussion of the relevant times). After incubation, the beads are imaged by epifluorescence microscopy. The bead fluorescence increases with the concentration of VEGF-A in the drop and saturates for concentrations larger than 4 × 10^4^ pg/ml, see Figure 2C-D. The sigmoidal shape of the calibration curve is typical of immunoassays [26]. The dynamical range of the immunoassay is given by the linear part of the curve, corresponding to concentrations of VEGF-A in the range 10^3^−4×10^4^pg/ml. The number of VEGF-A molecules in the droplet of volume *V*_*d*_ ≈ 50 nL is then in the range 7.5 × 10^5^ − 3.0 × 10^7^ molecules.

Let us compare this result with standard results obtained in 96-well plates. The range of detection advertised by the supplier for experiments in 96-well plates lies between 1 pg/mL and 2.3 ng/mL. The total number of VEGF-A molecules in the 50 *µ*L assay is then in the range 1.6 × 10^6^ − 1.8 × 10^9^ molecules. Note however that the number of beads in such an assay is around 20 times larger than in the microdroplet assays. The upper bound of the standard assay, i.e. the number of molecules leading to saturation, is therefore consistent with the experiments in microdroplets.

The standard assay has a lower detection limit than the microfluidic protocol for two reasons: (i) in the 96-well plate protocol, bead fluorescence is excited using a laser, which is more powerful than the epifluorescence lamp of our microscopy setup; and (ii) in the standard protocol, fluorescence is read with a photomultiplier, which typically has a better signal to noise ratio than our optical setup. The differences between the range of concentrations detectable in a standard 96-well plate assay and the microfluidic platform are summarized in **Table I**.

**Table I.** Adapting a 96-wells plate test to microfluidics. Between classical 96-well plates tests and our in-drop measurements, differences of bead number and sample volume are expected to increase the levels of concentration to which the test is sensitive

|  | Classic measurements | In-drop measurements |
| --- | --- | --- |
| Beads per assay | $\approx 100$ | $\approx 5$ |
| Sample volume | 50 μl | 50nl |
| Incubation time | 2x2h | 15h |
| Recommended VEGF-A concentration (pg/mL) | [ $1 \times 10^0 - 2.3 \times 10^3$ ] | [ $1.10^3 - 4.10^4$ ] |

### C. Quantifying the secretion of VEGF-A at the single spheroid level

We now use the platform to determine the amount of VEGF-A secreted by hundreds of individual spheroids on a single chip. After droplet production, spheroids are let to incubate for 4 hours, a time during which cells aggregate, the spheroid assembles, compacts, and reaches its circular steady-state shape (Supp. Fig. S1). The bead fluorescence is measured after 15 hours, and translated to an amount of secreted VEGF-A via the calibration curve shown in Figure 2D. The resulting distribution of extracellular VEGF-A is shown in Figure 3A. The histogram is not gaussian and skewed towards high concentrations of VEGF-A, with a mean concentration of VEGF-A of 2.4 ng/mL.

The same protocol is repeated on different chips, letting the spheroids incubate 24 or 48 hours before introducing the beads. Histograms of the concentration of secreted VEGF-A after 24 and 48 hours are presented in Figure 3B and C, respectively. As time increases, the mean concentration of extracellular VEGF-A increases, roughly doubling with every day of incubation, see Figure 3D. These experiments allow to calculate the secretion rate A of VEGF-A, giving A ≈ 15 − 20 molecules*/*spheroid*/*s. The distribution of the VEGF-A concentration widens with time (Figure 3A-C). Note that, while the distribution widens with time, the coefficient of variation of the distribution, defined as the ratio of standard deviation to the mean, remains constant: C*V* ≈ 40%, see Figure 3E.

**Figure 3.**
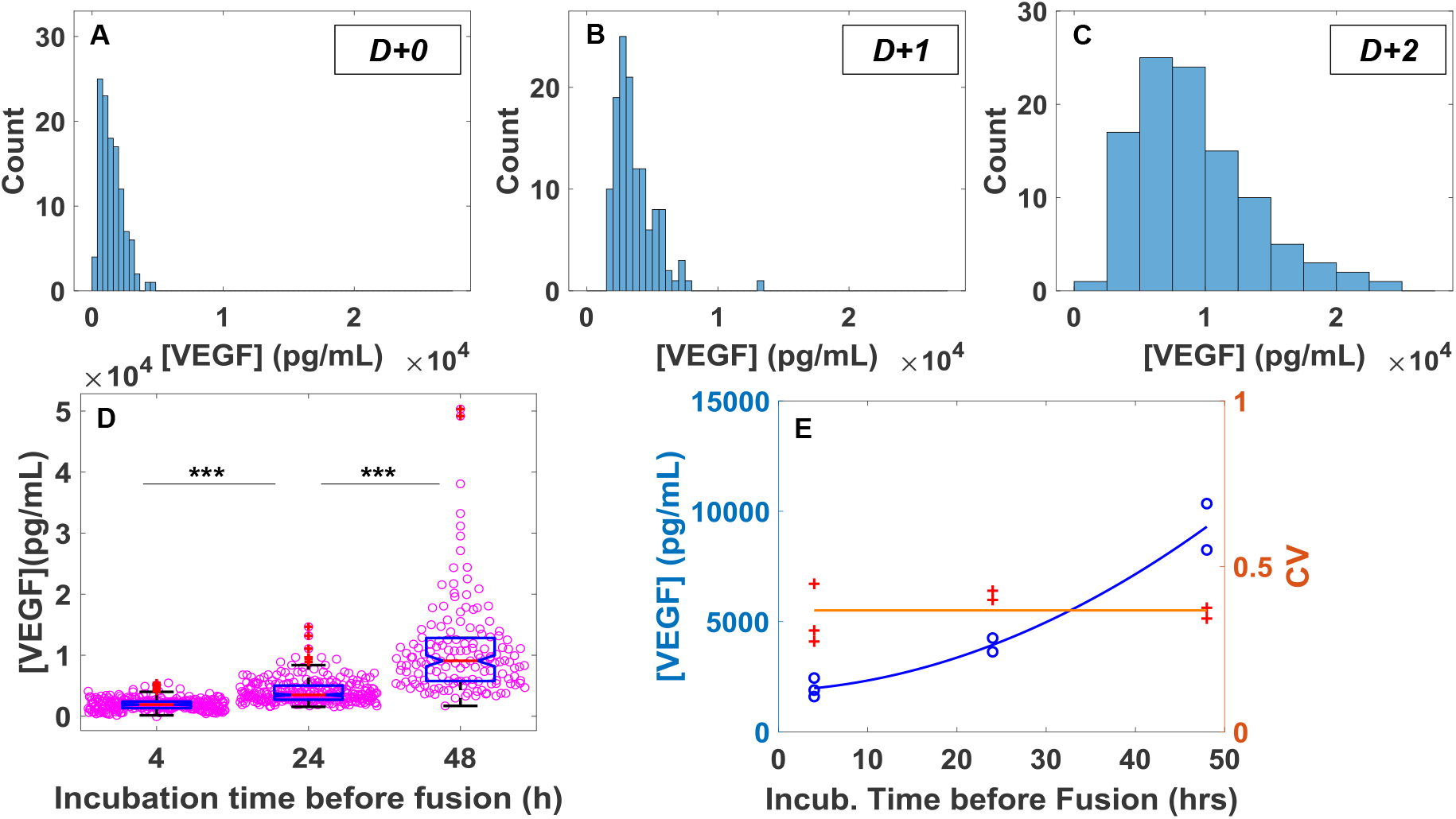
Quantification of the time evolution of VEGF-A secretion at single spheroid level. (A-C) Typical VEGF-A concentration distributions for individual chips where the spheroids were incubated for 4, 24, or 48 hours before bead introduction (D) Overall evolution of the secretion over days for individual spheroids. 4 hours: N=3 chips, n = 288 traps ; 24 hours: N=2 chips, n = 264 traps ; 48 hours: N=2 chips, n = 151 traps. Wilcoxon-Mann-Whitney tests, *** p<0.001. (E) Evolution of the mean and C.V of fluorescence distribution over time. Lines are shown to guide the eye.

When histograms are renormalized by their mean and standard deviations, they collapse onto each other and are well fitted by a Fréchet distribution of parameter *k, m* and *s*, with respective values (k=0.085;m=-7.5;s=7;), see Supp. Fig. S2. The Fréchet distribution can arise when the quantity observed, here the amount of VEGF-A secreted by one spheroid, is the result of a sum of underlying correlated random processes [32]. The correlation between the different processes may suggest differential regulation of the different steps of the protein secretions by individual spheroid (i.e. during the transcription, post-transcription, translation, post-translation, or exocytosis of the soluble protein).

### D. Combining measurements of cellular secretions on individual spheroids, and at the single-cell level

To understand the source of the heterogeneity in VEGF-A secretion, measurements are performed on the scale of each individual spheroid. This is done by using the present platform to combine measurements of the secreted molecule with a quantitative characterization of the spheroids. In particular we look for correlations between the secretions with the spheroid size and with the the intracellular concentration of VEGF-A within them.

We begin by investigating the influence of the spheroid size on VEGF-A secretion. In each drop, the spheroids are stained with a nuclear dye and imaged in wide-field epifluorescence microscopy to obtain the shape descriptors, as shown in Figure 4(A-C). In the particular chip analyzed in Figure 4, the distribution of radii is peaked around a mean value of 50 *µ*m, and displays a standard deviation of 12.5 *µ*m (Figure 4D). The extracellular VEGF-A concentration is quantified as previously, and the amount of VEGF-A produced by a single spheroid is plotted as a function of the spheroid projected area in Figure 4E, showing no correlation between the size of the spheroid and the amount of VEGF-A secreted.

**Figure 4.**
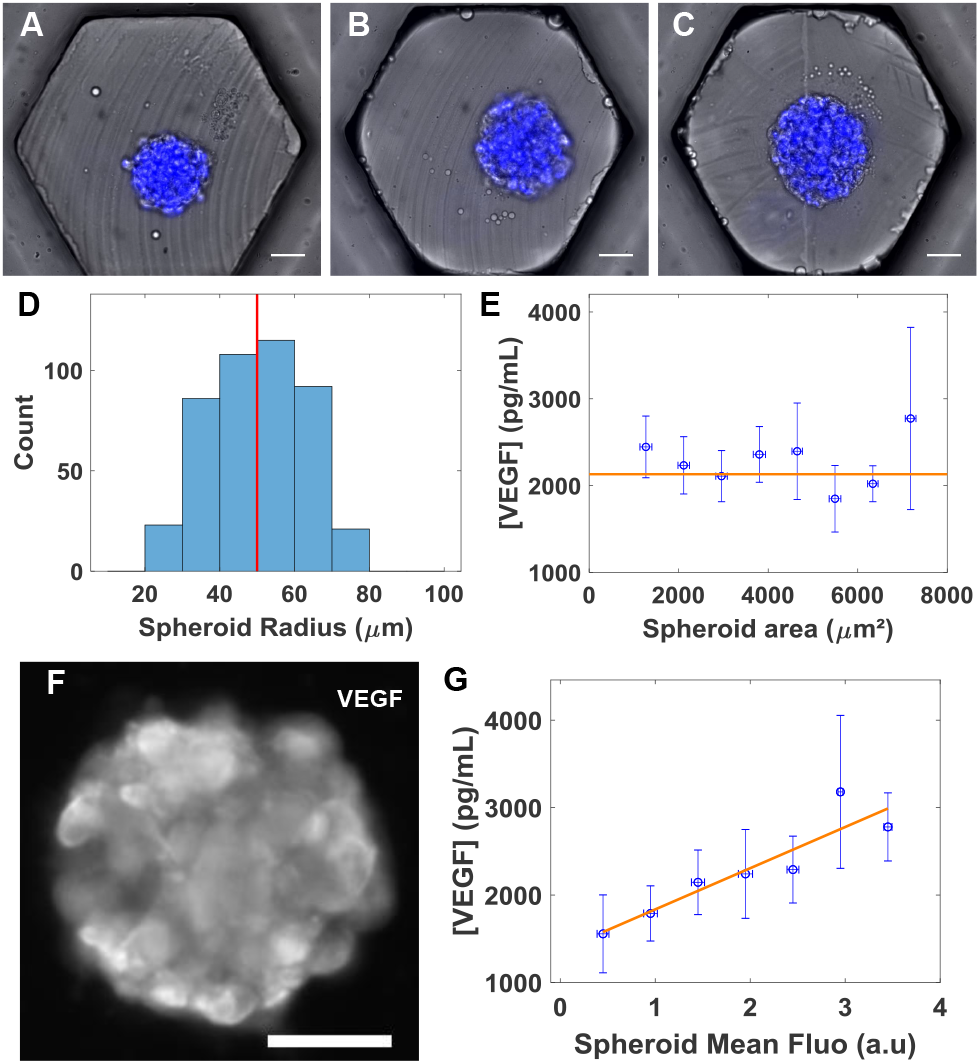
Correlation of the morphology and intracellular VEGF-A production with the secretion of the spheroids. (A-C) Spheroids are stained with a nuclear dye (blue) and imaged by epifluorescence microscopy. Scale bar: 50 *µ*m. (D) Distribution of spheroid radii within a single microfluidic device. Mean radius: 49.9 *µ*m, CV=24%. (E) Measurements of the secreted VEGF-A as a function of spheroid projected area measured by image analysis. All measurements are performed at D+1 after spheroid formation. The horizontal line is the average VEGF-A production. (F) Fluorescence image of a spheroid with VEGF-A immuno-staining. The production of VEGF-A is higher for cells at the periphery than in the center, see also [21]. (G) Correlation between VEGF-A secretions measured on a capture bead, and the mean intracellular amount of VEGF-A in a single spheroid, measured by ICC. The line is a linear fit (*R*^2^ = 0.843).

After completion of the bead immunoassay, the spheroids are fixed and stained by immunocytochemistry to detect the intracellular VEGF-A, following the protocols of Ref. [21]. A typical image of a spheroid after immunostaining is shown in Figure 4F, revealing a hetero-geneous distribution of VEGF-A within the spheroid. Indeed, Ref. [21] describes a detailed analysis of the signal within the spheroids through a layer-by-layer description, and shows that the production of VEGF-A is higher for cells at the periphery of the spheroid than in the center of the spheroid (see also Supp. Fig. S3). These measurements indicate that all cells do not secrete the same quantity of VEGF-A.

We now compare the extracellular concentration of VEGF-A to the average intracellular fluorescence within each spheroid. The resulting plot shows a clear correlation between intracellular production and amount of secreted proteins: spheroids exhibiting a higher level of intracellular fluorescence also secrete more VEGF-A, as shown Figure 4G. The heterogeneity in VEGF-A production is therefore not due to a size variability, but to the heterogeneity within the cell population, and the organization of these cells within each spheroid (See section IV).

### E. Modelling the molecular dynamics within droplets

To gain insight into the protocol described above and help the design of future experiments of secretion measurements, we turn to the modeling of our experimental system: an encapsulated spheroid secretes VEGF-A, which diffuses and eventually binds to a capture bead. The model involves several geometric and physical ingredients, which in turn can be summarized into three time scales that need to be accounted for: the secretion rate of VEGF-A, the time that a secreted molecule needs to reach the bead, and the rate of binding, which depends on the antibody affinity and on the local concentration of VEGF-A. It is important to understand the interplay between these time scales to interpret the signal on the bead at any given moment and to ensure that the immunoassay allows us to quantify the amount of cellular secretions.

We first o bserve t he effects of confinement on th e concentration wi thin th e droplet. For this, consider a system consisting of a spheroid of radius *R*_*s*_ = 50 *µ*m placed in a spherical droplet of radius *R*_*d*_ = 200 *µ*m, as sketched in Figure 5A. The spheroid secretes VEGF-A at a rate A, defined a s t he n umber o f m olecules r eleased by the s pheroid per unit time. The concentration of VEGF-A in the droplet is calculated using a simple diffusion model for the concentration *c*,

**Figure 5.**
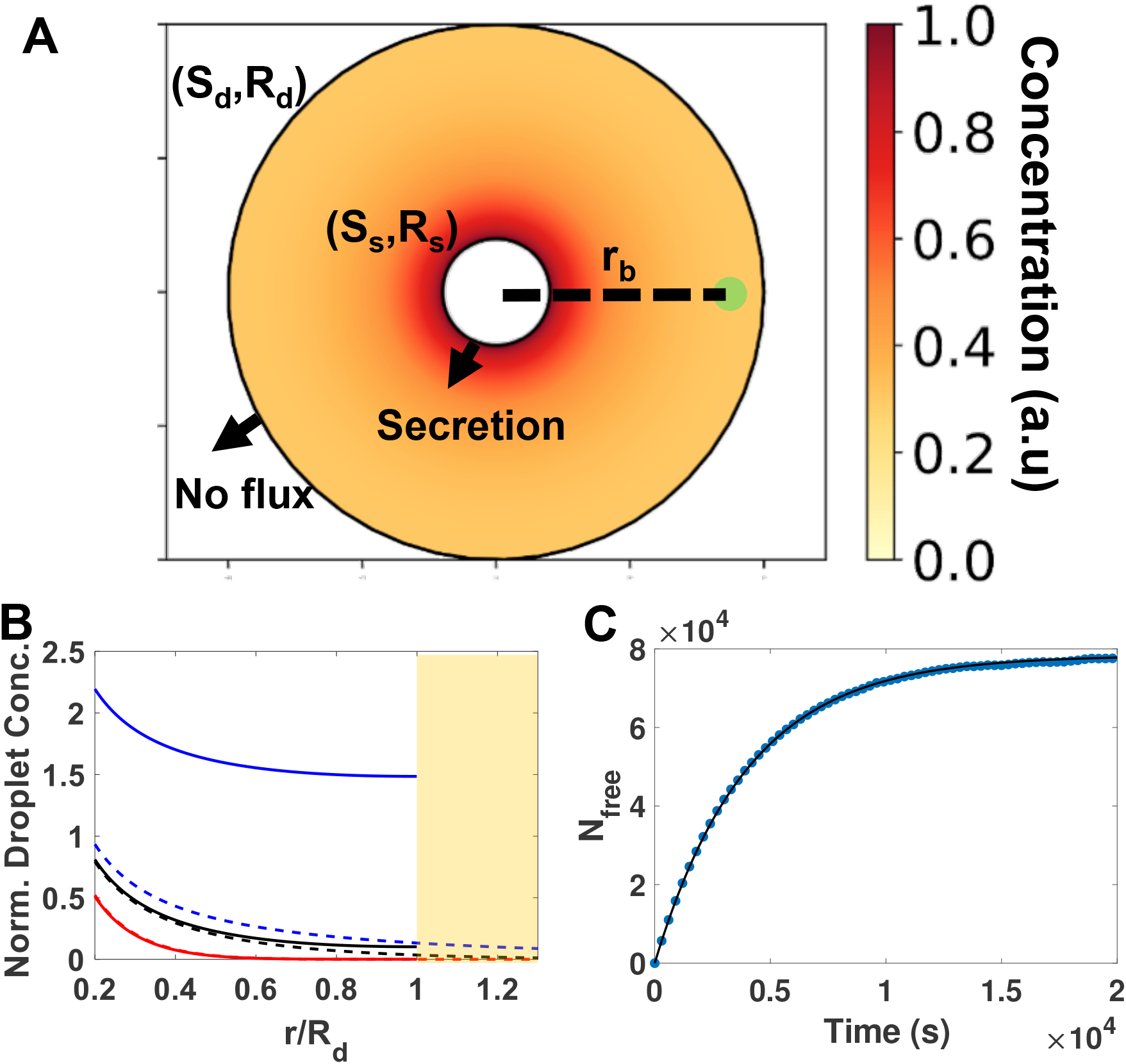
Modelling the dynamics of VEGF-A concentration within a confined droplet. (A) Schematic of the theoretical model of the drop (big circle), containing a centered spheroid (white circle) and a bead (green dot), colored as a function of concentration profile. (B) Concentration profile for VEGF-A in the droplet, with (solid) or without (dashed) confinement after times *t* = 0.9*τ*_*D*_ (red), 9.7*τ*_*D*_ (black) and 40*τ*_*D*_ (blue). In the unconfined case, the concentration profile converges to a stationary shape (dotted blue line). Concentration values are normalized by the maximum unconfined concentration value at steady-state. Secretion rate is 𝒜 = 96 molecules/s. Yellow color represents the outside of the droplet. (C) Smoldyn simulation (blue circles) of the number of VEGF-A molecules in the drop as a function of time, for a droplet of radius *R*_*d*_ = 200 *µ*m. The number of VEGF-A molecules increases before reaching a plateau. Its time evolution is well fitted by a saturating exponential (black line): 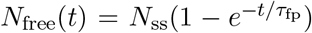, where 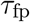is the mean first passage time.

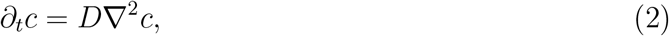

where *D* is the diffusion coefficient. The diffusing molecule is subject to the boundary condition

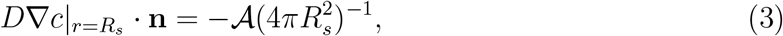

at the spheroid surface, where **n** indicates the outward facing normal to the surfaces. The far away boundary condition for the unconfined case is that *c*|_*r→∞*_ = 0, while the confined case has a no-flux boundary condition at the droplet surface:

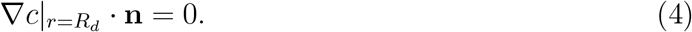

The first effect of confinement is that secretions accumulate over time in the droplet, contrary to the unconfined situation. This is visible in Figure 5B, where the concentration of VEGF-A is plotted in the confined and unconfined cases for three different times. Here time is made dimensionless by rescaling by the diffusion time within the droplet, 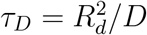. Figure 5B shows that the concentration of VEGF-A rises in a similar way in the confined and unconfined cases, until the molecules reach the edge of the droplet at *t* ≈ *τ*_*D*_. For our experimental parameters, the concentrations remain close even for *t* ≈ 10*τ*_*D*_. For larger times however, the concentration in the unconfined case remains nearly unchanged, while the concentration in the confined case increases on average within the droplet. For *t* = 40*τ*_*D*_ for instance, the local concentration at the surface of the spheroid is 2.5 times higher in the confined case, and is nearly 10 times higher at the droplet edge. This accumulation of the secreted molecule in the droplet makes its detection much easier in practice.

A more significant effect of the confinement is that all secreted molecules will eventually reach a capture bead placed in the droplet. This is in contrast to the case of a spheroid and capture bead placed in free space, where only a fraction of the secreted molecules find the bead [33]. To evaluate the time necessary for the molecules to reach the target, consider a capture bead of radius *a*, placed at a distance *r*_*b*_ away from the spheroid, and call *V*_*d*_ the droplet volume. The average time needed for a molecule of VEGF-A secreted by the spheroid to hit the bead is given by the mean first-passage time 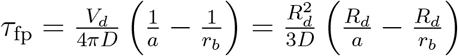 (see equation (50) in Ref. [34]). In our experiments, we have *r*_*b*_ ≈ *R*_*d*_, and *R*_*d*_≫*a*, so that 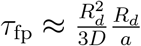. The time needed to find the bead is approximately given by the diffusion time needed to travel through the droplet 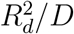, multiplied by the ratio of droplet size to target size *R*_*d*_*/a*: the smaller the area to hit, the more thoroughly the random walk needs to explore the domain before hitting the bead. Taking *D* = 1.33.10^*−*10^ m^2^/s for the diffusion coefficient of VEGF-A [35] and *a* = 5 *µ*m, we estimate *τ*_*D*_ ≈ 300 s and 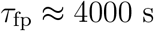.

The two other characteristic time scales can be estimated as follows: The secretion rate of the spheroid and the capture efficiency on the bead. The measured secretion rate of the spheroid is 𝒜 ≈ 20 molecules*/*s, see Figure 3, so that the associated time scale is *τ*_*A*_ = 1*/*𝒜 ≈ 0.05 s. The number of available capture sites *N*_*M*_ on the bead is *N*_*M*_ ≈ 1.5×10^7^, see Section III B. The corresponding concentration is 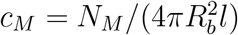, where *l* ≈ 30 nm is the size of an antibody. A typical rate for antigen-antibody binding is *k*_*on*_ = 10^6^ M^*−*1^.s^*−*1^ = 10^*−*18^ molecules^*−*1^.m^3^.s^*−*1^. The associated time scale is *τ*_*k*_ = 1*/*(*k*_*on*_ * *c*_*M*_) ≈ 10^*−*7^ s. We are therefore in a situation where 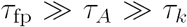: once a molecule is secreted by the spheroid, it takes a long time to find the capture bead, but then binds to it instantly.

We simulate the experimental system numerically with the software Smoldyn [36]. Smoldyn builds on a description of molecular reactions introduced by von Smoluchowski [28] to perform stochastic spatial simulations, where molecules are treated as point-like particles that diffuse and can react with each other or with surfaces. In a first simulation, a spheroid of radius *R*_*s*_ = 40 *µ*m is centered in a droplet of radius *R*_*d*_ = 200 *µ*m and secretes VEGF-A at a rate A = 20 molecules*/*s. The simulated capture bead has a radius *a* = 5 *µ*m and binding to it is considered to be diffusion-limited to avoid simulating 10^7^ binding sites on this curved surface, which would require much smaller simulation time steps [29, 37]. The diffusion coefficient of VEGF-A is taken to be *D* = 1.33 × 10^*−*10^ m^2^*/*s.

The simulated time evolution of the number of free VEGF-A molecules in the droplet is shown by the blue points in Figure 5C. The simulated data are well fitted by a saturating exponential *N*_free_(*t*) = *N*_ss_(1 − *e*^*−t/τ*^), where *N*_ss_ ≈ 7.83 × 10^4^ molecules and *τ* ≈ 4.00 × 10^3^ s (black line). The saturating exponential comes from the fact that we are counting the total number of VEGF-A molecules in the droplet over time. Molecules of VEGF-A are produced at a constant rate 𝒜, and captured at a rate 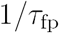, so that the equation of evolution for *N* is 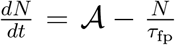, which solves as 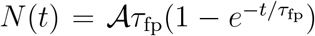.Note that the fit values are in excellent agreement with the theoretical values: the theoretical value for 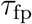 is 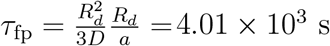, and the theoretical value for *N*_ss_ is 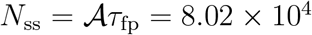molecules, within 3% of the fit value.

The simulation is repeated for three droplet radii: *R*_*d*_ = 100, 150 and 200 *µ*m, to highlight the influence of the droplet size on the first-passage time and therefore on the duration of the assay. The evolution of the number of free VEGF-A molecules in these droplets is plotted as a function of time in Figure 6A. In all cases, the simulated data are well fitted by a saturating exponential 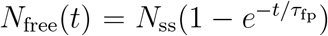, with excellent agreement between the fit values and theoretical values: decreasing the droplet radius decreases the mean first passage time and the steady-state number of free VEGF-A molecules in a droplet proportionally to the droplet volume. For a droplet of radius *R*_*d*_ = 100 *µ*m (resp. 150 *µ*m), the fitted first passage time is 516 s (resp. 1681 s), to compare to the theoretical value of 501 s (resp. 1692 s). The numerical steady-state values of the number of free VEGF-A molecules in solution are likewise in very good agreement with the theoretical values 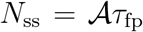, with less than 5% discrepancy between fit and theory. Fitted values are *N*_ss_ = 1.04×10^4^ molecules for a droplet of radius *R*_*d*_ = 100 *µ*m and *N*_ss_ = 3.21 × 10^4^ molecules for a droplet of radius *R*_*d*_ = 150 *µ*m. These simulations highlight the critical influence of the droplet radius on the duration of the assay. The mean first passage time 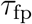is proportional to the volume of the droplet, so that doubling the droplet radius leads to an eight-fold increase in the mean first passage time 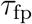.

**Figure 6.**
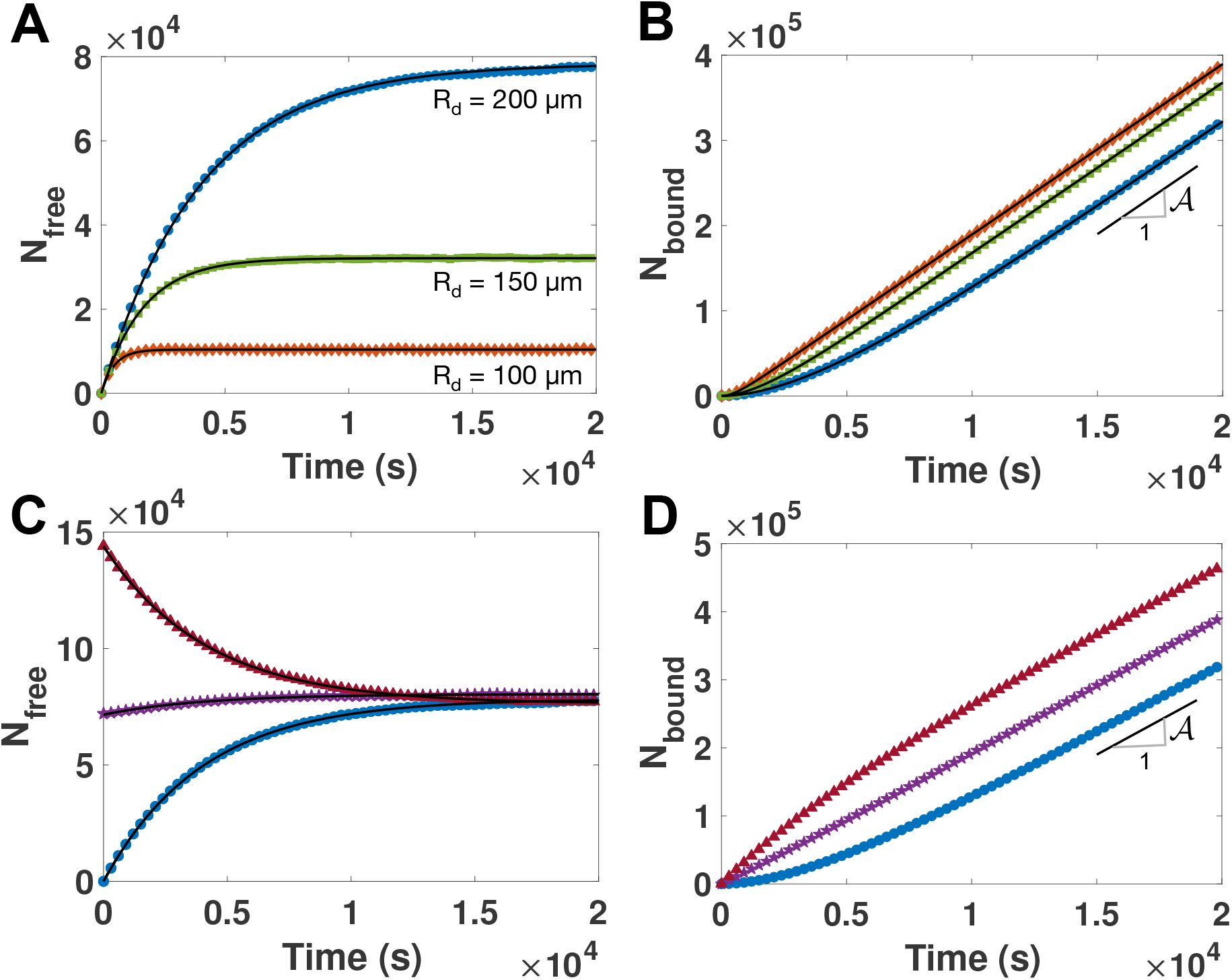
Simulated binding dynamics in droplets. (A) Simulated evolution of the number of free VEGF-A molecules in the droplet as a function of time, for three droplet radii. Orange diamonds: *R*_*d*_ = 100 *µ*m, green squares: *R*_*d*_ = 150 *µ*m, blue circles: *R*_*d*_ = 200 *µ*m. Black lines are fit to 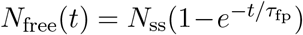.Fitted values of 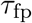: 516 s, 1681 s and 4004 s. Associated saturation values: 10381, 32111 and 78313 molecules. (B) Time evolution of the number of bound molecules on the bead. Simulations (points) and theoretical shape (black lines): *N*_bound_(*t*) = 𝒜*t* − *N*_free_(*t*). Same color legend as in (A). The black line segment has a slope 𝒜 = 20 molecules*/*s. (C) Effect of pre-incubation on the number of free VEGF-A molecules in a droplet of radius *R*_*d*_ = 200 *µ*m. Capture bead introduced without incubation (blue circles), with an incubation time of 1 hour (purple stars) or 2 hours (red triangles). These molecules bind to the bead in a time ≈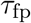, and then the steady-state regime where capture balances secretion is attained. (D) Number of bound molecules on the bead for the three incubation times, showing the quick initial capture of available molecules, and then the linear evolution of the number of bound molecules with time. The black line segment has a slope 𝒜 = 20 molecules*/*s.

The experimental readout is the number of bound molecules *N*_bound_ on the capture bead, which also increases with time, see points in Figure 6B. The evolution of the number of bound molecules is given by 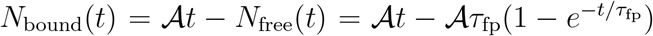, see black lines in Figure 6B. After a transient time of order 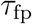, the number of bound molecules increases linearly with time: for 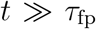, we have *N*_bound_(*t*) ≈ 𝒜*t*. Therefore, at any given time, the number of bound molecules is larger in smaller droplets because the mean first-passage time is smaller, but after a transient 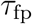, the rate of increase of the number of bound molecules is A, independent of the droplet size. Note that the total time needed to saturate the bead is 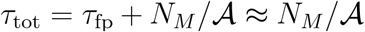 for 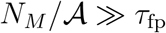. In our experiments, using *N*_*M*_ = 10^7^ binding sites and A = 20 molecules.s^*−*1^, the time needed to saturate the bead is ≈ 6 days, much longer than our experiments.

Finally, note that the secretions of spheroids in the experiments above are measured after the spheroid is incubated in the droplet for a time *τ*_*i*_, before introducing the capture bead. One can therefore ask how long it takes for the already secreted molecules to reach the bead, and how the incubation time influences the duration of the assay. We simulate pre-incubation times *τ*_*i*_ = 1 and 2 hours. The number of free VEGF-A molecules in solution when the capture bead is introduced is 𝒜*τ*_*i*_. These molecules also diffuse and bind to the capture bead in a characterestic time 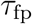, during which the spheroid continues to secrete. After a transient regime, secretion is again balanced by capture and the number of free VEGF-A molecules in the bulk is constant up to stochastic fluctuations, see Figure 6C. On the capture bead, the number of bound molecules increases quickly over the first 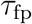 seconds, after which the kinetics of binding go back to their steady-state dynamics, see Figure 6D. The time to saturate the bead is therefore *τ*_tot_ ≈ (*N*_*M*_ − 𝒜*τ*_*i*_)*/*𝒜, considering as before that 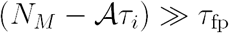.

To summarize, the limiting time scale in our problem is the first-passage time 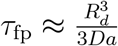, which is proportional to the droplet volume and inversely proportional to the radius of the capture bead. In our experimental setup, 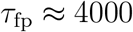, and measurements of the number of captured VEGF-A molecules on the bead make sense only for 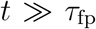, which rationalizes our choice of letting the bead incubate for 15 hours before measuring its fluorescence.

## IV.DISCUSSION AND CONCLUSIONS

The present study demonstrates how anchored microfluidic droplets can be used to form hundreds of spheroids and how the droplet confinement within an oil phase can facilitate the accumulation and measurement the secreted molecules in the aqueous solution. These capabilities are achieved by leveraging the newly developed asymmetric anchors [25], which allow us to add functionalized beads to the droplet containing each spheroid, at any time after the spheroid formation. Fluorescence measurements on these beads then allow us to quantify the secretome, at any time during the culture (Fig. 1).

We apply this technique to measure the production of VEGF-A, one of the important molecules secreted by hMSCs. While the secretion of VEGF plays a key role *in vivo* to promote angiogenesis and wound healing, its expression is inhibited when the hMSCs are cultured in standard 2D formats. Working in the 3D format is therefore fundamental for promoting VEGF-A secretion. Using our microfluidic assay, we find a secretion rate of ≈ 20 VEGF molecules/spheroid/s. The rate is preserved for different pre-incubation times of 4, 24 or 48 hours, indicating that the spheroids secrete at a relatively constant rate over the two days of culture.

Moreover, our measurements on 100-250 spheroids in parallel evidence the inherent heterogeneity in the production of VEGF-A by the *mesenchymal bodies* (MBs). This heterogeneity, which had not been observed previously, does not correlate with the heterogeneity in spheroid size. Instead, the ability to link intracellular and extracellular secretions of VEGF-A at the spheroid level confirms that the variations in the secreted VEGF-A are well correlated with intra-cellular measurements. This sheds new light on the level of functional diversity at the scale of each of the individual spheroids. Indeed, our previous measurements on intra-cellular VEGF-A showed a wide variety of production levels VEGF-A that were linked, on the single-cell level, with the differentiated status of each of these progenitor cells [21]. In turn, the differentiation state dictates the cell organization in 3D, as the cells form coherent MBs, in which the production of VEGF-A is strong on the edge of the MBs and weak in the central region. Taken together, the new results confirm that VEGF expression is regulated through the interactions of individual cells within each spheroid, and that the spheroid thus acts as a functional cellular unit [21].

Finally, the theoretical model that we introduce rationalizes the performance of droplet-based immuno-assays, based on time-scales describing the behavior of soluble molecules within the droplets and on the capture beads. Confinement is found to play a major role, first by allowing the molecules to accumulate in the droplets, then by ensuring that all of the molecules eventually find their way to the measurement beads. It is important to realize that the time to reach the bead, given by the first-passage time 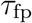, increases linearly with the volume of the droplet, and that readouts of the number of bound molecules on the capture bead make sense only when 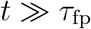. To keep measurement times reasonables, the droplet size for microfluidic immunoassays should therefore result from a compromise between being a large enough reservoir of nutrients for the spheroid, while still being small enough for capture to occur within a few hours, which is the typical time scale of immuno-assays.

Looking ahead, this new microfluidic platform will enable new experiments that combine spheroid culture, stimulation, and measurements of intra-cellular and secreted molecules. The ability to link the soluble molecules with the cell fate will be important to understand mechanisms of cell-cell interactions, e.g. for immune-cancer interactions, as well as providing a marker for cell response to drugs, e.g. for pharemaceutical screening. And as advanced data methods become widespread, the large data sets provided by these experiments will provide new ways to address complex cellular processes.

## ACKNOWLEDGEMENTS

We gratefully thank Caroline Frot for her support on microfabrication, Tharshana Stephen from the Cytometry and Biomarkers Unit of Technology and Service (CB UTechS) for her help with cytometry tools, and Steve Andrews for help with Smoldyn. We also acknowledge the friendly help of the Image Analysis Hub of the Institut Pasteur for this work. This project has received funding from the European Research Council (ERC) under the European Union’s FP7 research and innovation program under Grant Agreement 714027.

## SUPPLEMENTARY FIGURES

**Figure S1.**
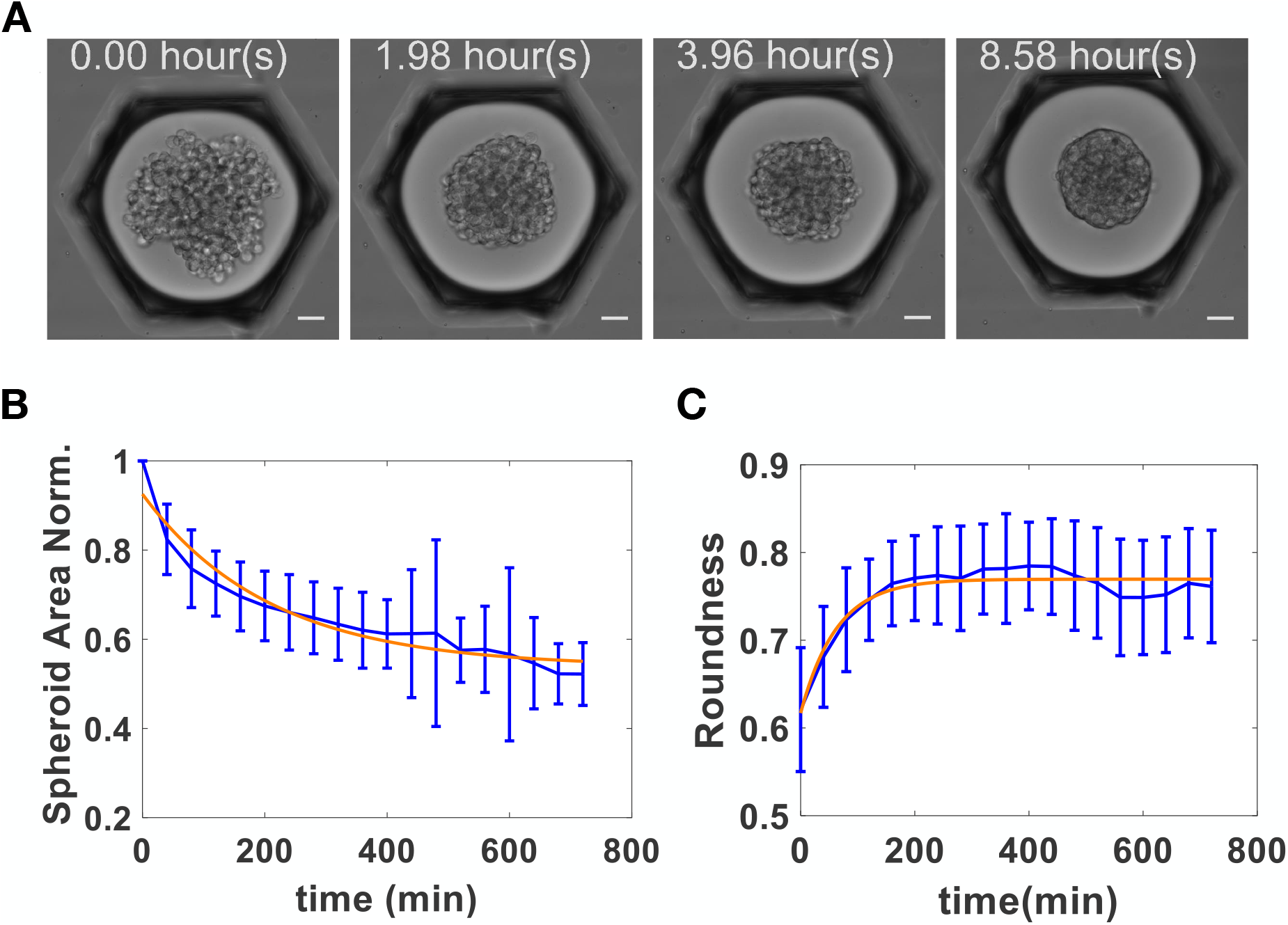
Spheroid formation. (A) Time lapse of spheroid formation and compaction in an anchored microfluidic droplet. (B) Average spheroid area as a function of time, renormalized by the spheroid area at initial time. Error bars show the standard deviation. (C) Average spheroid roundness as a function of time. Calling *S* the spheroid area and *P* its perimeter, roundness is defined as *S/*(4*πP* ^2^).

**Figure S2.**
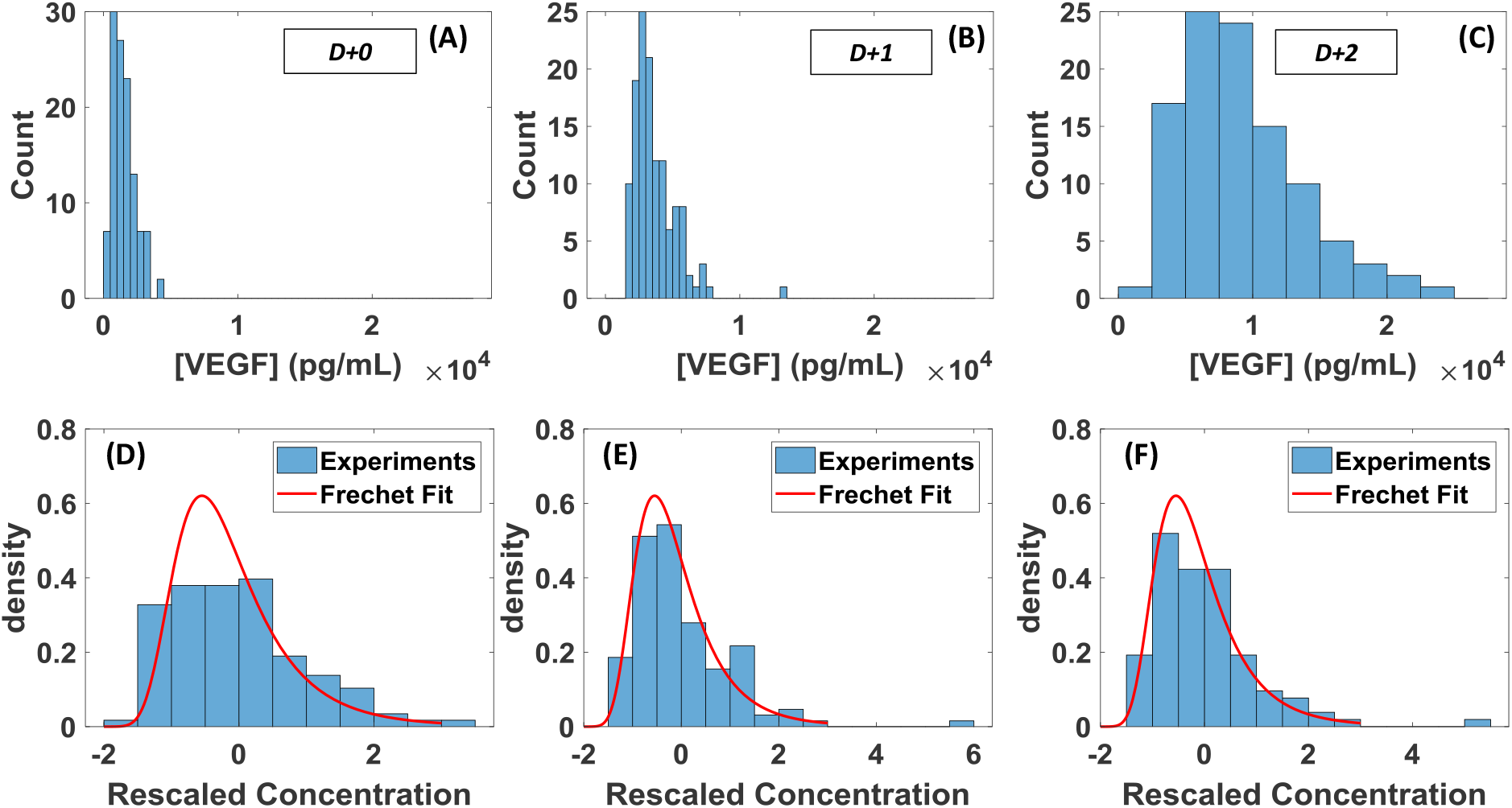
Rescaled histograms of secreted VEGF-A. Top row: histograms of secreted VEGF-A. Bottom row: same histograms, rescaled by subtracting the mean and dividing by the standard deviation, and Fréchet fit: 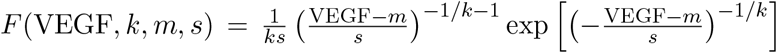with *k* − 0.085, *m* = −7.5, *s* = 7. The same parameter values also fit data on the distribution of proteins in a population of bacteria [32]. Data taken at (A) D+0, (B) D+1, (C) D+2.

**Figure S3.**
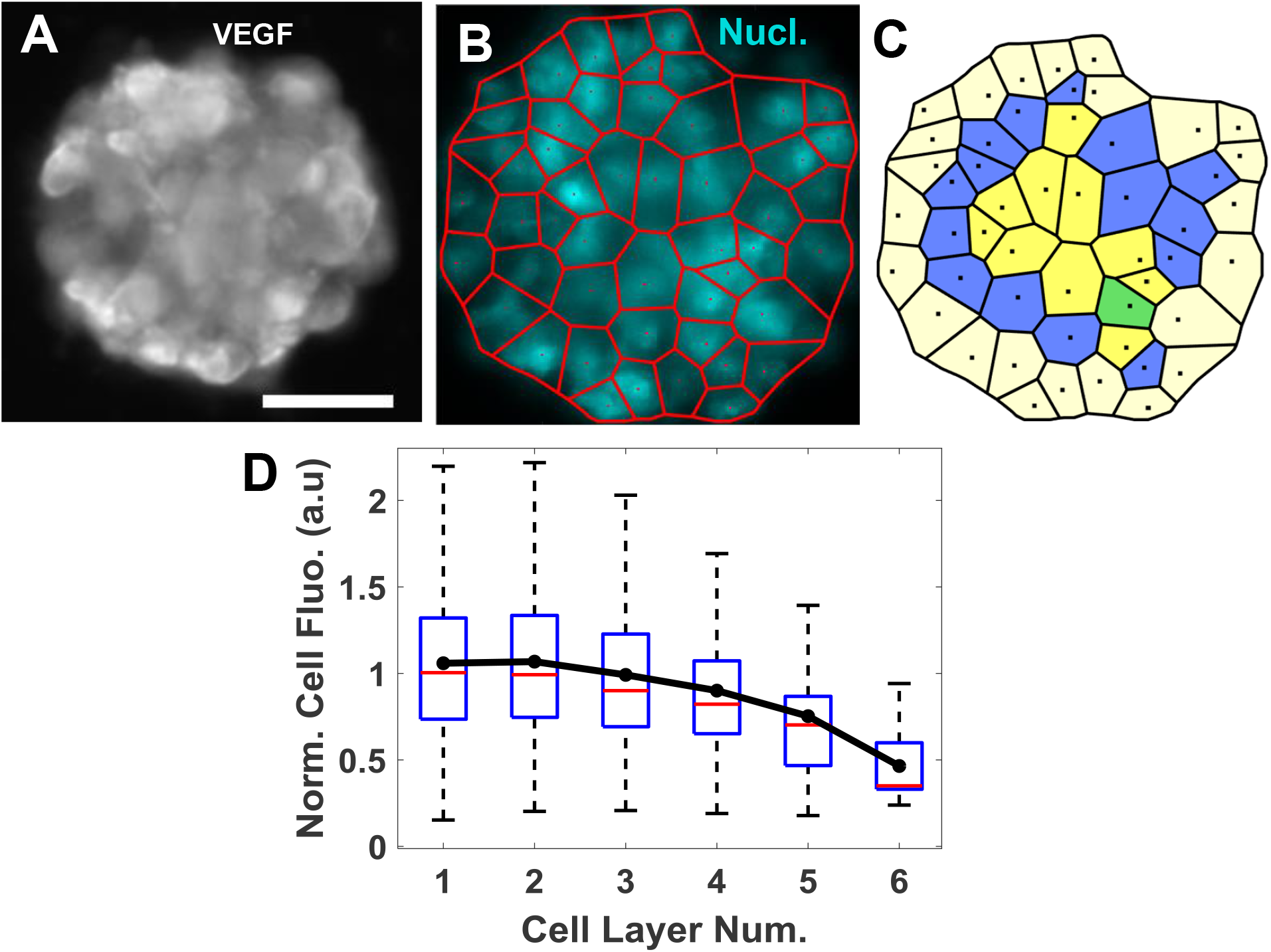
Layer-by-layer analysis of VEGF production in spheroids. (A) Epifluorescence microscopy image of a spheroid stained by ICC for VEGF-A. (B) Epifluorescence microscopy image of cell nuclei stained with DAPI, for the same spheroid as in (A). The centroids of nuclei are identified (red dots), and individual cells are defined by a Voronoi tesselation from the centroids (red lines). (C) Definition of cell layers in the spheroid. Pale yellow: first layer, blue: second layer, bright yellow: third layer, green: fourth layer. (D) VEGF-A fluorescence signal as a function of cell layer, showing an increased VEGF secretion in the outer layer compared to the core. The VEGF-A signal is renormalized by its average value in the outer layer. Error bars show the standard deviation.

